# ViruSurf: an integrated database to investigate viral sequences

**DOI:** 10.1101/2020.08.10.244624

**Authors:** Arif Canakoglu, Pietro Pinoli, Anna Bernasconi, Tommaso Alfonsi, Damianos P. Melidis, Stefano Ceri

## Abstract

ViruSurf, available at **http://gmql.eu/virusurf/**, is a large public database of viral sequences and integrated and curated metadata from heterogeneous sources (GenBank, COG-UK and NMDC); it also exposes computed nucleotide and amino acid variants, called from original sequences. A GISAID-specific ViruSurf database, available at **http://gmql.eu/virusurf_gisaid/**, offers a subset of these functionalities. Given the current pandemic outbreak, SARS-CoV-2 data are collected from the four sources; but ViruSurf contains other virus species harmful to humans, including SARS-CoV, MERS-CoV, Ebola, and Dengue.

The database is centered on sequences, described from their biological, technological, and organizational dimensions. In addition, the analytical dimension characterizes the sequence in terms of its annotations and variants. The web interface enables expressing complex search queries in a simple way; arbitrary search queries can freely combine conditions on attributes from the four dimensions, extracting the resulting sequences.

Several example queries on the database confirm and possibly improve results from recent research papers; results can be recomputed over time and upon selected populations. Effective search over large and curated sequence data may enable faster responses to future threats that could arise from new viruses.

## INTRODUCTION

The pandemic outbreak of the coronavirus disease COVID-19, caused by the virus species SARS-CoV-2, has created unprecedented attention towards the genetic mechanisms of viruses. The sudden outbreak has also shown that the research community is generally unprepared to face pandemic crises in a number of aspects, including well-organized databases and search systems. We respond to such urgent need by means of a novel integrated database and search system collecting and curating virus sequences with their properties. Data is captured, standardized, organized, and made accessible to the scientific community, so as to facilitate current and future research studies.

In our work, we are driven by the Viral Conceptual Model (VCM) for virus sequences (1), which was recently developed by interviewing a variety of experts of the various aspects of virus research (including clinicians, epidemiologists, drug and vaccine developers). The conceptual model is general and applies to any virus. The sequence of the virus is the central information; sequences are analyzed from a *biological dimension* describing the virus species and the host environment, a *technological dimension* describing the sequencing technology, an *organizational dimension* describing the project which was responsible for producing the sequence, and an *analytical dimension* describing properties of the sequence, such as known annotations and variants. Annotations include known genes, coding and untranslated regions, and so on. Variants are extracted by performing data analysis and include both nucleotide variants – with respect to the reference sequence for the specific species – with their impact, and amino acid variants related to the genes.

We have previously proposed another conceptual model focused on human genomics (2), which was based on a central entity representing files of genomic regions, similarly described from various dimensions. We next developed and implemented an integrated database (3), searchable through the GenoSurf (5) interface (http://gmql.eu/genosurf/). Thanks to such previous knowledge in human genomics, we have been able to rapidly design VCM and then to deploy ViruSurf.

Currently, ViruSurf includes sequences from GenBank (17) of SARS-CoV-2 and SARS-related coronavirus, as well as MERS-CoV, Ebola and Dengue viruses; the pipeline is generic and other virus species will be progressively added next, giving precedence to those species which are most harmful to humans. For what concerns SARS-CoV-2, we also include sequences from COG-UK (21) and NMDC (http://nmdc.cn/). GenBank and COG-UK data are made publicly available and can be freely downloaded and re-distributed. Special arrangements have been agreed with GISAID (8; 19), resulting in a GISAID-enabled version of ViruSurf. Due to constraints imposed by GISAID, the database exposed in this version lacks the original sequences, certain metadata and nucleotide variants; moreover, GISAID requires their dataset not to be merged with other datasets. Hence, the two versions of ViruSurf should be used separately, and a certain amount of integration effort must be carried out by the user.

Although the origin sources provide well-organized data portals (NCBI Virus (11), the COVID-19 Data Portal https://www.covid19dataportal.org/, and GISAID EpiCoV data browser), these do not allow for an integrated search over multiple sources, nor they provide fast selection using sequence variants. A number of other integrated interfaces are being developed in alternative research contexts: UCSC SARS-CoV-2 Genome Browser (9); 2019nCoVR (24) at the Chinese National Genomics Data Center; VirusDIP (22) at the China National GeneBank; CovSeq (13); CARD (18). Compared to other resources, ViruSurf has much stronger query and search capabilities; we use the power of conceptual modeling to structure metadata and to organize data integration and curation; we support search queries allowing to combine filters on metadata, nucleotide and amino acid mutations in an effective and scalable way, treating all of them as first-class citizens. The closest comparison is with CoV-GLUE (20) (only containing GISAID data), which includes variants, but forces users to search one variant at a time, visualizing the list of sequences with that variant.

## MATERIALS AND METHODS

### Database Content

For SARS-CoV-2, ViruSurf contains data from four main data sources: NCBI (including Genbank and RefSeq), COG-UK, NMDC, and GISAID. We also reviewed other available sources (GenomeWarehouse and CNGdb) but observed that they do not add substantial value to the integration effort, as most of their sequences overlap with those stored in the four cited sources.

Table 1 provides a description of the current ViruSurf content: for each virus we report the rank, ID and name from NCBI Taxonomy, the number of sequences included from each source and the reference sequence. We next provide the average number per sequence of each annotation and nucleotide/amino acid variants computed against the reference sequence. Note that, although GISAID uses a different reference sequence, provided amino acid variants are relative to protein sequences (which are the same as in other sources), hence they can be compared with other variants.

**Table 1.** Summary of ViruSurf content as of August 4th, 2020. For each taxon name (identified by a taxon ID and rank) and each source, we specify the number of distinct sequences and the reference genome; we also provide the average number of annotations, nucleotide variants and amino acid variants per sequence. The GISAID-only entry refers to those GISAID sequences that are not also present in the other three sources.

| Taxon rank | Taxon ID | Taxon name | Source | #Seq. | Reference | Avg Annot. | Avg Nuc. Var. | Avg AA Var. |
| --- | --- | --- | --- | --- | --- | --- | --- | --- |
| No rank | 2697049 | Severe acute respiratory syndrome coronavirus 2 | GISAID all | 76,664 | EPI_ISL_402124 | - | - | 4.8 |
| No rank | 2697049 | Severe acute respiratory syndrome coronavirus 2 | GISAID only | 46,366 | EPI_ISL_402124 | - | - | 4.7 |
| No rank | 2697049 | Severe acute respiratory syndrome coronavirus 2 | GenBank + RefSeq | 13,309 | NC_045512.2 | 28.0 | 17.6 | 24.0 |
| No rank | 2697049 | Severe acute respiratory syndrome coronavirus 2 | COG-UK | 38,124 | NC_045512.2 | 28.0 | 25.5 | 58.0 |
| No rank | 2697049 | Severe acute respiratory syndrome coronavirus 2 | NMDC | 295 | NC_045512.2 | 28.1 | 28.8 | 56.5 |
| Species | 694009 | Severe acute respiratory syndrome-related coronavirus | GenBank + RefSeq | 673 | NC_004718.3 | 14.0 | 91.6 | 19.7 |
| Species | 1335626 | Middle East respiratory syndrome-related coronavirus | GenBank + RefSeq | 1,381 | NC_019843.3 | 27.0 | 104.0 | 87.4 |
| Species | 2010960 | Bombali ebolavirus | GenBank + RefSeq | 5 | NC_039345.1 | 9.0 | 126.8 | 41.6 |
| Species | 565995 | Bundibugyo ebolavirus | GenBank + RefSeq | 22 | NC_014373.1 | 9.0 | 130.5 | 40.9 |
| Species | 186539 | Reston ebolavirus | GenBank + RefSeq | 57 | NC_004161.1 | 9.0 | 126.1 | 29.4 |
| Species | 186540 | Sudan ebolavirus | GenBank + RefSeq | 40 | NC_006432.1 | 9.0 | 368.6 | 51.0 |
| Species | 186541 | Tai Forest ebolavirus | GenBank + RefSeq | 9 | NC_014372.1 | 9.0 | 4.6 | 2.8 |
| Species | 186538 | Zaire ebolavirus | GenBank + RefSeq | 2,938 | NC_002549.1 | 9.0 | 503.7 | 66.3 |
| Strain | 11053 | Dengue virus 1 | GenBank + RefSeq | 11,185 | NC_001477.1 | 15.0 | 469.7 | 200.9 |
| Strain | 11060 | Dengue virus 2 | GenBank + RefSeq | 8,692 | NC_001474.2 | 15.0 | 410.0 | 117.7 |
| Strain | 11069 | Dengue virus 3 | GenBank + RefSeq | 5,344 | NC_001475.2 | 15.0 | 269.3 | 118.6 |
| Strain | 11070 | Dengue virus 4 | GenBank + RefSeq | 2,492 | NC_002640.1 | 15.0 | 265.1 | 147.2 |

Some content of ViruSurf is extracted from the sources and used without changes, some is manually curated, and some is computed in-house (nucleotide and amino acid variants, impact, quality measures). The code repository is available on GitHub (https://github.com/acanakoglu/virusurf downloader). The current content corresponds to data available at the sources on August 4th, 2020. We will provide periodical updates on a monthly basis.

### Relational Schema

The core schema, represented in Figure 1, is inspired to classic data marts (4), with a central fact table describing the Sequence, featuring several characterizing attributes, and then four dimensions:

**Figure 1.**
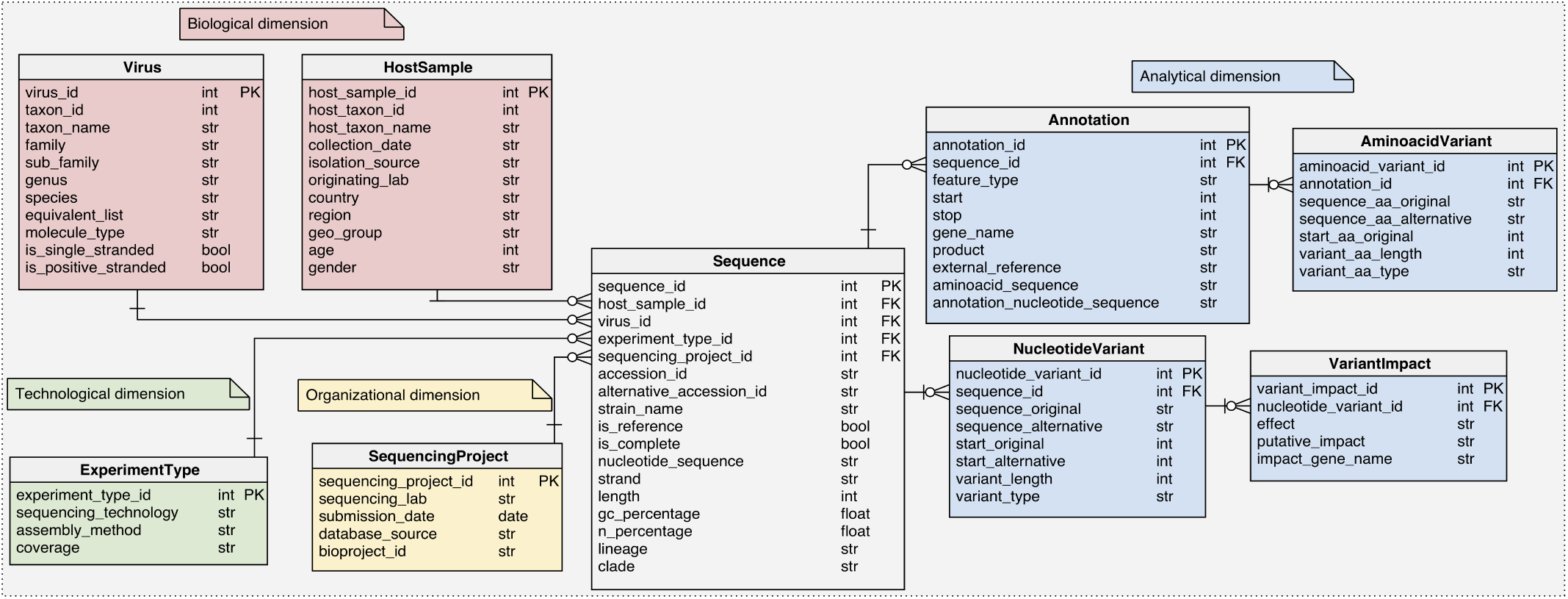
Logical schema of the relational database in the back-end of ViruSurf.

1. The *Biological dimension* (including the VIRUS and HostSample tables) is concerned with the virus species characterization and the host organism, including the temporal/spatial information regarding the extraction of the biological sample.
2. The *Technological dimension* (ExperimentType table) describes the sequencing method.
3. The *Organizational dimension* (SequencingProject table) describes the project producing each sequence.
4. The *Analytical dimension* provides annotations for specific sub-sequences and characterizes the variants in the nucleotide sequence and in the amino acid sequence. It includes the Annotation, AminoacidVariant, NucleotideVariant and VariantImpact.

All tables have a numerical sequential primary key (PK), conventionally named <`table name`*>* `id` and indicated as PK in Figure 1; we indicate with foreign keys (FK) the relationships from a non-key attribute to a primary key attribute of a different table. Relationships from the Sequence towards Virus, HostSample, SequencingProject and ExperimentType are functional (e.g. one Sequence has one ExperimentType, while an ExperimentType may be the same for multiple Sequence); instead, relationships in the analytical dimension are 1:N (e.g. one Sequence has many Annotations, and an Annotation has many AminoacidVariants). In the Supplementary Table S1 we describe the specific attributes of every table.

### Data Import

The pipeline used to import the content of the ViruSurf database from sources is shown in Figure 2. We use different download protocols for each source:

**Figure 2.**
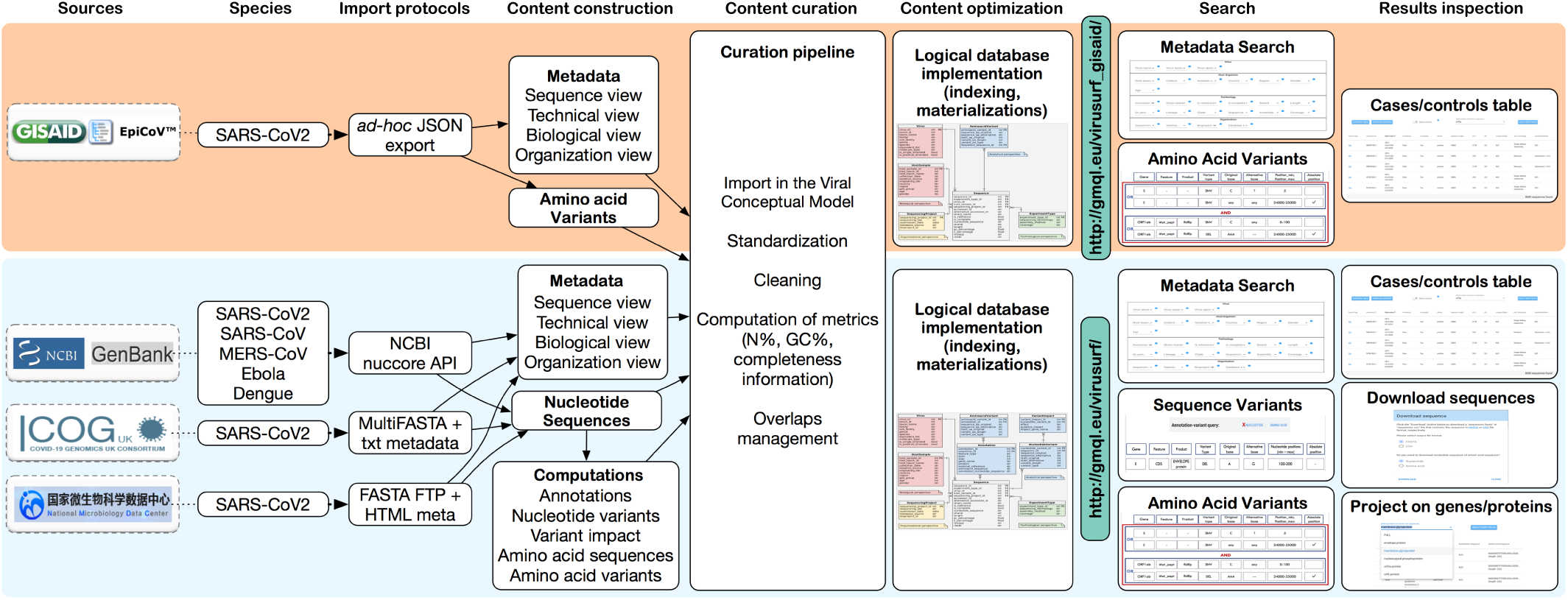
General pipeline of the ViruSurf platform. For given sources and species, we use download procedures to construct content, perform data curation, and load the content into two distinct databases, for GISAID and for the other sources, which are schema-compatible (the former is a subset of the latter). We then provide two Web-based interfaces supporting search and result inspection.

- For NCBI data (including GenBank and RefSeq sequences), we employ the extraction tools available in the E-utilities (16): the Python APIs allows to retrieve one complex XML file for each sequence ID available in NCBI.
- COG-UK instead provides a single MultiFASTA file on its website (https://www.cogconsortium.uk/data/); this is associated with a text file for metadata.
- NMDC exposes an FTP server with FASTA files for each sequence, while metadata are captured directly from the HTML description pages.
- GISAID provides to us an export file in JSON format, updated every 15 minutes. The file is produced by GISAID technical team in an *ad-hoc* agreed form for ViruSurf.

Automatic pipelines have been implemented to extract metadata and fill the Sequence, Virus, HostSample, SequencingProject, and ExperimentType tables; some attributes require data curation, as next described.

### Annotation and Variant Calling

In order to provide homogeneous information for sequence annotations and variants, we use a unique annotation procedure for GenBank, COG-UK and NMDC; resulting variants for amino acid sequences are consistent with those provided by GISAID. We extract: structural annotations, nucleotide and amino acid sequences for each annotated segment, nucleotide variants and their impact, amino acid variants for the proteins, and other information such as percentage of specific nucleotide bases.

For each virus, we manually select a reference sequence and a set of annotations, comprising coordinates for codifying and structural regions, as well as the amino acid sequences of each protein. Usually, such data are taken from the RefSeq entry for the given virus (e.g., NC 045512 for SARS-CoV-2). For each imported sequence, the pipeline starts by computing the optimal global alignment to the reference by means of the dynamic programming Needleman-Wunsch (NW) algorithm (14). The time and space complexity of NW is quadratic in the length of the aligned sequences, which often hinders its adoption in genomics, but viral sequences are relatively short, thus we can use NW rather than faster heuristic methods. We configured the algorithm to use an affine gap penalty, so as to favor longer gaps which are very frequent at the ends of sequences.

Once the alignment is computed, all the differences from the reference sequence are collected in the form of variants (substitutions, insertions or deletions). Using the SnpEff tool (7) we annotate each variant and predict its impact on the codifying regions; indeed, a variant may, for example, be irrelevant (e.g, when the mutated codon codifies for the same amino acid of the original codon), produce small changes, or be deleterious. Based on the alignment result, the sub-sequences corresponding to the reference annotations are identified within the input sequence.

Coding regions are then translated into their equivalent amino acid sequences; the translation takes into consideration annotated ribosomial frameshifts events (e.g., within the ORF1ab gene of SARS-CoV-2). When translation fails (e.g., because the nucleotide sequence retrieved from the alignment is empty or its length is not a multiple of 3), we ignore the amino acid product; failures are due to incompleteness and poor quality of the input sequence, further computation of amino acid variants would produce erroneous information. Instead, when an aligned codon contains any IUPAC character ambiguously representing a set of bases (https://genome.ucsc.edu/goldenPath/help/iupac.html), it is translated into the *X* (unknown) amino acid, which automatically becomes a variant. Note that queries selecting known amino acids are not impacted; unknown amino acids are usually not of interest. Translated amino acid sequences are then aligned with the corresponding amino acid sequences (using NW), annotated with the reference, and amino acid variants are inferred.

Alignment, variant calling, and variant impact algorithms are computationally expensive, so we decided to parallelize this part of the pipeline, taking advantage of Amazon Elastic Compute Cloud (Amazon EC2). We implemented a chunked and parametrized execution modality for distributing the analysis of the sequences associated to each virus to multiple machines, so that the total execution time of the process can be divided by the number of available machines.

### Data Curation

Our first curation contribution is to provide a unique schema (VCM (1)) for different data sources. Metadata of each source use different terms, the mappings between VCM attribute names and those used at the original sources are in Supplementary Table S2. Specific value curation efforts have been dedicated to the location information, the collection and submission dates, the completion of virus and host taxonomy names/identifiers, the choice of the appropriate reference sequence (cross-checking with several research papers to ascertain that the typical reference sequence used for variant calling is defined), the coverage of the sequencing assay. We also compute metrics regarding the percentage of G and C bases, of unknown bases and the information about sequence completeness.

Some sequences are deposited to multiple sources; we detect such redundancy by matching sequences based on their strain name or the pair of strain name and length. Overlaps among sources is illustrated in Figure 3. As all overlaps occur between GISAID and the other sources, we store the information about overlaps within the GISAID database and allow the possibility of performing “GISAID only” queries, i.e., restricted to GISAID sequences that are not present in GenBank, COG-UK or NMDC.

**Figure 3.**
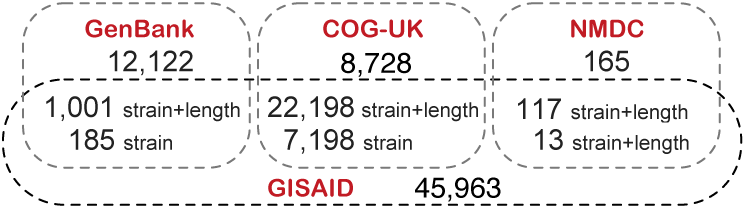
Counts of SARS-CoV-2 overlapping sequences from each source. Overlaps are computed by means of either the strain name, or both strain name and length.

## RESULTS

### Web Interface

The web interface of ViruSurf is composed of 4 sections, numbered in Figure 4: (1) the menu bar, for accessing services, documentation and query utilities; (2) the search interface over metadata attributes; (3) the search interface over annotations and nucleotide/amino acid variants; (4) the result visualization section, showing resulting sequences with their metadata. Results produced by queries on the search interface (2) are updated to reflect each additional search conditions, and counts of matching sequences are dynamically displayed to help users in assessing if query results match their intents. The interface allows to choose multiple values for each attribute at the same time (these are considered in *disjunction*); it enables the interplay between the searches performed within parts (2) and (3), thereby allowing to build complex queries given as the logical conjunction – of arbitrary length – of filters set in parts (2) and (3).

**Figure 4.**
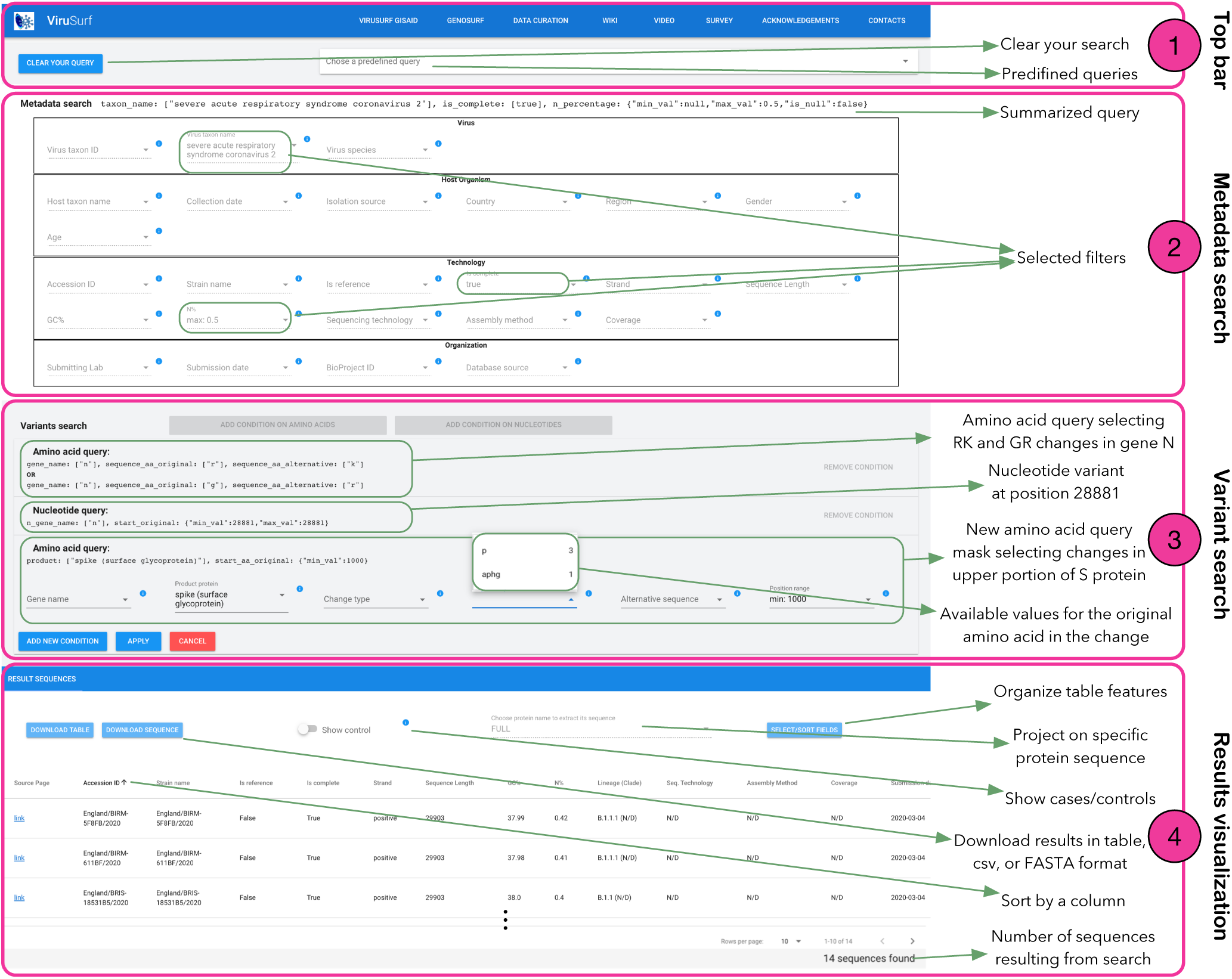
Overview of ViruSurf interface with example of data query combining three filters on metadata (Part 2) and three variant query panels (in Part 3): one amino acid query and one nucleotide query are already applied; one amino acid query is open and in the process of completion. In Part 4, 3 resulting sequences already reflect the choices applied in Parts 2 and 3.

#### Menu bar

The menu bar includes links to the GISAID-specific ViruSurf system, to the GenoSurf system, and pointers to the data curation detail page, to the wiki, to a video compilation, and to a pedagogical survey supporting the user by documenting the aspects of search queries; on the top right of the interface we provide various ‘Predefined queries’.

#### Metadata Search

The Metadata search section is organized in four parts: *Virus* and *Host Organism* (from the *biological* dimension), *Tecnology* and *Organization* (from the respective dimensions). It includes attributes which are present in most of the sources, described by an information tab that is opened by clicking on blue circles; values can be selected using drop-down menus. At the side of each value we report the number of items in the repository with that value.

The user can compose desired queries by entering values from all the drop-down menus; the result is the set of sequences matching all the filters. Note that the special value N/D (Not Defined) indicates the null value, that can also be used for selecting items. For numerical fields (age, length, GC% and N%) the user must specify a range between a minimum and maximum value; in addition, the user can check the N/D flag, thereby including in the result those sequences having the value set to N/D. Similarly, collection date and submission date have a calendar-like drop-down components, supporting a range of dates and the N/D flag.

#### Variant Search

The Variant search section allows searching sequences based on their nucleotide variants (with their impact) and the amino acid variants. When the user selects ‘ADD CONDITION ON AMINO ACIDS’ or ‘ADD CONDITION ON NUCLEOTIDES’ buttons, a dedicated panel is opened, with a series of drop-down menus for building search conditions. A user can add multiple search conditions within the same panel; these are considered in disjunction. Once the panel is completed, it is registered; registered panels can be then deleted from a query, if needed. Variants selected in different panels are intended in conjuction. Heterogeneous variant searches (i.e., on amino acid and nucleotide ones) can only be combined in panels, thus in conjunction.

In the example shown in Figure 4 (which represents the construction of the ‘Predefined query 8’, from Pachetti et al. (15)) the user is choosing all SARS-CoV-2 sequences that are complete, have a maximum percentage of unknown bases of 0.5%, and have R to K `or` G to K amino acid changes in gene N `and` a nucleotide variant at position 28,881. The filters set up to this point have selected a set of 14 sequences (as indicated at the bottom right of the page) In the represented snapshot, a third variant panel is in the process of being compiled with an amino acid condition that could be added to the two existing ones by pressing ‘APPLY’. This holds a filter on the spike protein and on the position of the variant on the protein (*>* 1000); the filter on original amino acid allows to select P (3 sequences available) or APHG (1 sequence).

#### Result Visualization and Download

The result table describes the sequences resulting from the selections of the user. The columns of the table can be ordered/included/excluded from the visualization; the resulting table can be downloaded for further processing. Whenever the user either adds or removes a value in the Metadata search, by clicking on a drop-down menu, the results table is updated; instead, it is updated only when a panel of the Variant search section is complete.

The ‘Show control’ switch allows to visualize the sequences of the control group, defined by those sequences selected by the Metadata search filters for which there exist some variant and the variant filters are not satisfied. This option, suggested to us by virologists, is the most sensible for describing the effects of variant analysis.

A user can select from a drop-down menu which sub-part of a nucleotide or amino acid sequence should be visualized; the default returns a ‘FULL’ nucleotide sequence (leaving the amino acid field empty), but with this menu option it is possible to return in the result the specific segment of interest. The whole result table – as it is visualized, inclusive of selected metadata and nucleotide or amino acid sequences – can be downloaded for further analysis as a CSV. Alternatively, the user may download either full or selected sequences by using their accession ID, either as CSV or FASTA files.

#### GISAID-specific ViruSurf

ViruSurf presents a version that is specific for data imported from GISAID, as requested by a specific Data Agreement. This interface presents limited functionalities but is nevertheless powerful and allows for combining its results with the ViruSurf main interface. Notable differences are here summarized: 1) After selecting filters, a user must explicitly apply her search by pressing an execution button. 2) Searches may be performed on the full dataset from GISAID or on the specific subset of sequences that are only present on GISAID (button ‘Apply GISAID specific’) – this may result particularly useful when the user wishes to compare or sum up results from the two interfaces (see Q3 in the following for an example). 3) Both drop-down menus and the result table’s columns hold the original GISAID attribute name, when available – when this differs from ViruSurf’s, the second one is provided in second position inside parentheses.

### Example queries

By means of complex search queries over our database it is possible to help virus research, according to the requirements provided by several domain experts; this is not currently supported by existing systems, which typically offer very nice visual interfaces reporting results of data analysis but limited search capabilities. We cite some examples inspired by recent research works.

**Q1**. To support SARS-CoV-2 vaccine design efforts, it is useful to track antigenic diversity. Typically, pathogen genetic diversity is categorized into distinct *clades* (i.e., a monophyletic group on a phylogenetic tree). In Gudbjartsson et al. (10), specific sequence variants are used to define clades/haplogroups (e.g., the “A3 group” is characterized by the 11083 and 29742 nucleotides G mutated to T, by the 1397 nucleotide G mutated to A, and by the 28688 nucleotide T mutated to C). ViruSurf supports all the information required to replicate the definition of SARS-CoV-2 clades proposed in the study.

**Q2**. In SARS-CoV-2, the G-T transversion at 26144, which caused an amino acid change in ORF3 protein (G251V), is investigated in Chaw et al. (6). The paper claims that this mutation showed up on 1/22/2020 and rapidly increased its frequency. We can use ViruSurf to find out that GenBank currently provides 3 complete sequences with such mutation collected before 1/22/2020, while GISAID provides other 13 sequences, non-overlapping with GenBank ones.

**Q3**. A study from Scripps Research, Florida, found that the mutation D614G stabilized the SARS-CoV-2 virus’s spike proteins, which emerge from the viral surface. As a result, the viruses with D614G seem to infect a cell more likely than viruses without that mutation; the G genotype was not present in February and was found with low frequency in March; instead, it increased rapidly from April onward. The scientific manuscript by Zhang et al. (23), cited by mass media (https://www.nytimes.com/2020/06/12/science/coronavirus-mutation-genetics-spike.html), has not been peer reviewed yet, but others on the same matter are (12)).

ViruSurf can be used to illustrate this trend. Let us consider two queries on complete sequences. Sequences with the D614G mutation collected before March 30 are 6,592, against 4,664 without the mutation; sequences with the D614G mutation collected after April 1 are 23,649, against 3,331 without the mutation. In both queries, case/control checks are obtained by using the ‘Show control’ switch, which retrieves – for the population specified by metadata filters – sequences that either have or do not have the chosen variants.

The same queries can be repeated on the GISAID-specific version of ViruSurf. Sequences with the D614G mutation collected before March 30 are 15,034, against 8,821 without the mutation; sequences with the D614G mutation collected after April 1 are 18,421, against 3,369 without the mutation. By summing up the query results from the two non-overlapping databases, we obtain that the sequences with the D614G mutation are 61.6% of those collected before March 30 and 86.3% of those collected after April 1.

## DISCUSSION

ViruSurf provides a single point of access to curated and integrated data resources about several Virus species. Example queries show that ViruSurf is able to replicate research results and to monitor how such results are confirmed over time and within different segments of available viral sequences, in a simple and effective way. The relevance of ViruSurf as a tool for assisting the research community will progressively increase with the growth of available sequences and of the knowledge about viruses.

While today’s efforts are concentrated on SARS-CoV-2, ViruSurf can similarly be useful for studying other virus species, such as other Coronavirus species, the Ebola Virus, the Dengue Virus and MERS-Cov, epidemics which are a current threat to mankind; ViruSurf will also enable faster responses to future threats that could arise from new viruses, informed by the knowledge extracted from existing virus sequences available worldwide.

In our future work, we plan to add epitopes (amino acid subsequences that can used for designing vaccines, with their lineage, host, evidence, and type of response), retrieved from IEDB and stored within new ViruSurf tables, and then searched using suitable Web interface extensions. We also plan to design data analysis services that can provide sophisticated use of our big data collection.

## ACKNOWLEDGEMENTS

The authors would like to thank Ilaria Capua, Matteo Chiara, Ana Conesa, Luca Ferretti, Alice Fusaro, Ruba Khalaf, Susanna Lamers, Stefania Leopardi, Francesca Mari, Carla Mavian, Graziano Pesole, Alessandra Renieri, Anna Sandionigi, Stephen Tsui, Limsoon Wong and Federico Zambelli for their contribution to requirements elicitation and for inspiring future developments of this research. The authors are grateful to the GISAID organization for the data sharing agreement that allowed the development of the GISAID-specific version of ViruSurf. The authors also acknowledge the depositions of worldwide laboratories to GenBank, COG-UK and NMDC. Finally, we acknowledge the support from Amazon Machine Learning Research Award “Data-driven Machine and Deep Learning for Genomics”.

